# Pore-scale hydrodynamics influence the spatial evolution of bacterial biofilms in a microfluidic porous network

**DOI:** 10.1101/436337

**Authors:** Jayde A. Aufrecht, Jason Fowlkes, Amber N. Bible, Jennifer Morrell-Falvey, Mitchel J. Doktycz, Scott T. Retterer

## Abstract

Bacteria occupy heterogeneous environments, attaching and growing within pores in materials, living hosts, and matrices like soil. Systems that permit high-resolution visualization of dynamic bacterial processes within the physical confines of a realistic and tractable porous media environment are rare. Here we use microfluidics to replicate the particle shape and packing density of natural sands in a 2D platform to study the flow-induced spatial evolution of bacterial biofilms underground. We discover that initial bacterial dispersal and particle attachment is a stochastic process driven by bacterial rheotactic transport across pore space velocity gradients. Over time, we find that gravity-driven flow conditions activate different cell-clustering phentoypes in EPS producing and EPS defective bacteria strains, which subsequently changes the overall spatial distribution of cells across the porous media network as colonies grow and alter the fluid dynamics of their microenvironment.

## Introduction

Microbial communities are complex systems that shape, and are shaped by, their local microenvironments. Bacteria often inhabit heterogeneous microenvironments with hydrodynamic flows that influence local nutrient transport and chemical gradients, creating specialized niches for microorganisms^1,2,3^. Likewise, the spatial confinement of some bacteria can influence emergent phenomena like quorum sensing, intercellular communication, and biofilm formation^4,5,6^. These microenvironment factors and complexities contribute to microbial community diversity and synergism, which hinders their isolation and culture in bulk laboratory conditions^7,8^.

Soil exemplifies a complex and heterogeneous microbial environment. Within soil, the physical and chemical structure of the porous network dictates water and nutrient flow, influencing the cells’ spatial distribution, communal behaviour, evolution, and even cross-kingdom interactions^9–11^. Some bacterial species can produce extracellular polymeric substances (EPS), which improve a soil’s moisture retention and act to aggregate soil particles together, further altering underground hydrodynamics^12,13^. Thus, bacterial characteristics are tightly coupled with the dynamics of soil conditions. This bacteria-soil interplay has implications for bioremediation, water quality, nutrient cycling, and underground ecology. It is therefore necessary to study soil bacteria within the structural and hydrodynamic context of their natural environment and on length scales appropriate to cellular functions (i.e. the pore scale) to elicit emergent behaviours. Unfortunately, the opacity of soil presents a challenge for the direct visualization and measurement of bacterial traits at the pore-scale *in situ*.

Experiments in sand columns and micromodels have enabled measurements of bulk bacterial transport through porous media and have even allowed some preliminary imaging of bacteria in pore-spaces^14–17^. Nafion, a transparent fluoropolymer that is sometimes used as a sand substitute, can further increase the imaging compatibility of bacteria in flow cells^18,19^. However, these systems do not have a defined structure and are often treated as ‘black boxes’, making it impossible to correlate pore-scale hydrodynamics with bacterial biofilm distribution.

In other synthetic systems, microfluidic platforms have been used to visualize bacterial behaviour in flow through narrow channels and around tight corners^20,21,22^. These platforms reduce the physicochemical complexity of natural porous media while testing bacterial characteristics in highly parameterized and fully defined systems^23^. Microfluidic systems have the added benefit of retaining the same physical structure for each experimental replicate, allowing flow in the channels to be computationally simulated^24,25^. These systems have elucidated bacterial chemotaxis through tortuous channels, bacterial streamer formation, and microbial competition^9,26^. However, there has yet to be a microfluidic system used to study bacterial transport and spatial distribution that replicates the heterogeneous structure of natural porous media.

Previous efforts have used microfluidic designs to re-create homogeneous porous features. Several such platforms rely on a simplistic design of circular features with varied packing densities, radii, and pitches^27–30^. While the reduction of particle shape to equivalent spheres has been successfully used in the field of soil physics to describe averaged fluid space quantities such as porosity and permeability, the size of a single bacterial cell is on a length scale well below the representative pore space volume that justifies the spherical particle assumption^31,32^. In order to inform individual based models of microbial community behaviours, an accurate particle shape is needed to reproduce the fluid dynamics and labyrinthine pore spaces that an individual microbe would experience underground.

Some microfluidic designs have taken into consideration the heterogeneity of natural porous media. Pore throat networks inspired by reservoir rock have been reconstructed through the use of Voronoi cell tessellations^33^, and focused ion beam-scanning electron microscopy (FIB-SEM) image triangulation, but geometrically partitioning the pore space results in a loss of information from the original sample^33,34^. Another team used a particle generator to create ellipsoid-shaped particles with a pore space reminiscent of natural granular media and, within their platform, demonstrated the influence of EPS on underground hydration processes^35^. The ellipsoidal shapes of the generated particles, however, are not based on shape statistics collected from natural grains.

In this work, we replicate both the shape and packing distributions of natural sand in a 2D microfluidic platform. A reduction in dimension from natural 3D soil substrates, while retaining the porosity of 3D soils, enables a quick image acquisition time to capture the dynamics of bacteria-particle interactions. Our heterogeneous design retains the pore-scale physical structure of natural porous media while providing a fully defined, tractable model for characterizing pore space hydrodynamics. Using gravity fed flow, we characterize the spatial evolution of bacterial biofilms in the porous network and correlate biofilm attachment and growth to pore scale hydrodynamics. We find that fluid shear trapping, resulting from rheotaxis, has the biggest influence on the initial spatial distribution of bacteria, but over time EPS production ability and biofilm expansion ultimately determines the spatial distribution of bacteria in this heterogeneous network.

## Results and Discussion

### Device design and simulation of pore space hydrodynamics

The microfluidic particle design was created using a granular media algorithm to randomly generate 500 particles from the size and shape distributions of natural sands^36,37^. Sand was chosen as a physical model for soil, the latter consisting of variable amounts of sand, silt, clay, and humus that form micro- and macro-aggregate structures. The resulting porous media section of the microfluidic platform contains a heterogeneous network of pore sizes and shapes including constriction points that are too small for cells to pass through to large pores with high connectivity (Figure 1, a-d). The distribution of pore spaces closely resembles a lognormal function with a median width of 50 μm (Supplementary Figure 1). A large variety of pore features allows for many sub-experiments on biofilm growth in various hydrodynamic conditions to be conducted at once.

**Figure 1:**
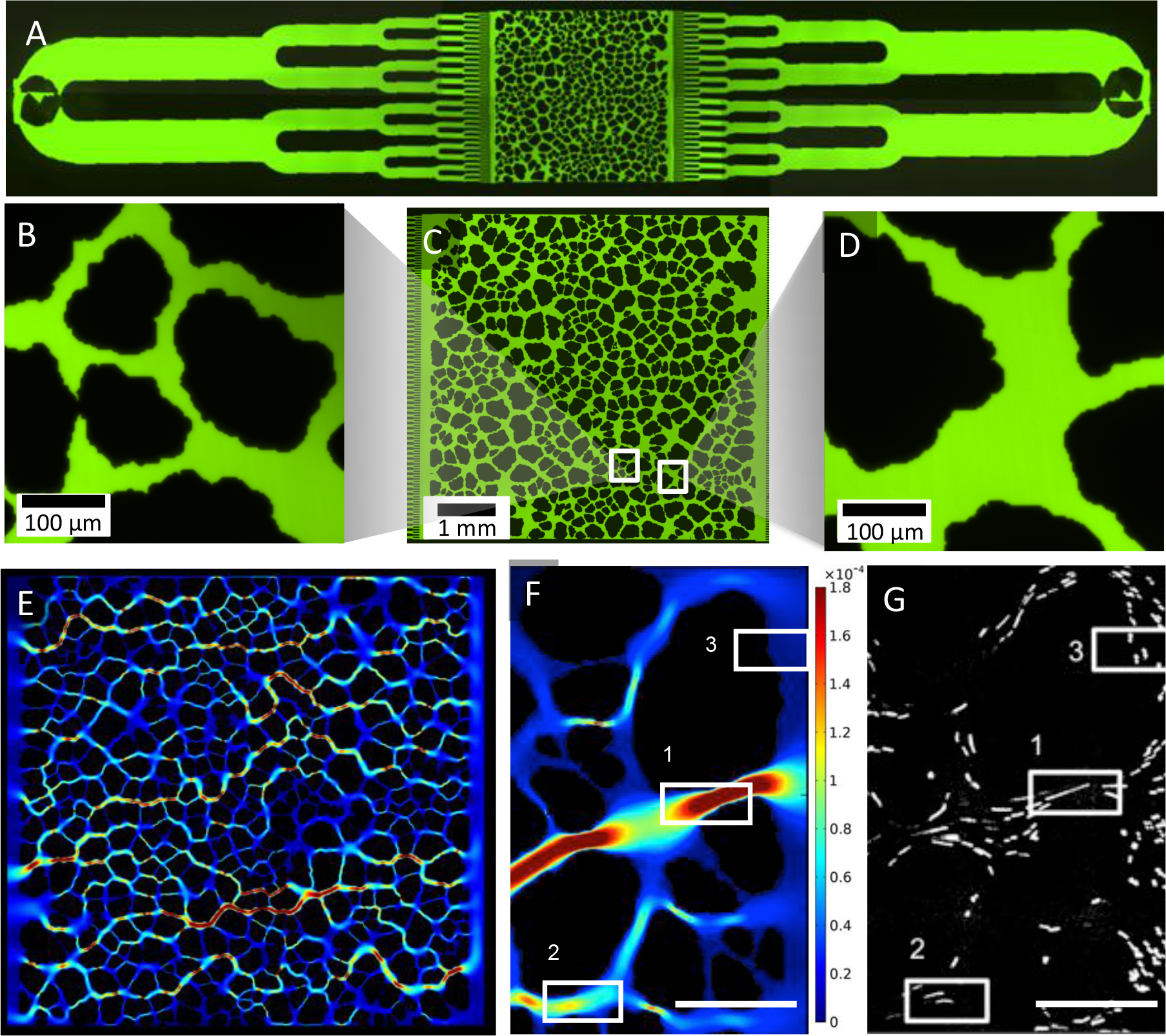
A microfluidic platform replicates the natural shape and layout of sand particles. (A) The device has a bifurcating inlet and outlet to uniformly distribute bacterial cells across the design (C) The porous network design (approximately 6,000 x 6,000 x 10 um) consists of heterogeneous pore spaces that have (B) constriction points smaller than the size of a bacteria cell and (D) larger, highly connected pore spaces. (E) Velocities within the pore space were simulated using COMSOL Multiphysics, which illuminated multiple preferential flow paths across the system. (F,G) The pore space velocities were experimentally verified using particle image velocimetry and velocity magnitudes (F, regions 1, 2, 3, decreasing in magnitude) corresponded to bead speeds within the same pore (G, regions 1, 2, and 3)(scale bars = 100 μm, velocity units = m/s)

The microfluidic porous media maintains the same particle layout and characteristics for each experimental replicate, which provides a highly tractable method to correlate pore scale features with flow. Here the CAD file used to fabricate the microfluidic platform also created the geometry of a COMSOL Multiphysics laminar flow simulation to determine the hydrodynamic flow parameters at steady state for constant velocity inputs. A velocity magnitude heat map was generated to visualize flow between pores and illuminated several preferential flow paths across the system (Figure 1, E). To confirm the accuracy of the COMSOL model, particle image velocimetry was used to calculate average flow speeds within pores, which matched the velocity magnitudes simulated in the COMSOL model (Figure 1, F-G).

### Bacteria growth and flow rate are coupled in a pressure driven system

To recreate the flow of percolating rainwater, a reservoir was suspended above the microfluidic platform and connected by tubing, generating gravity-driven flow. Prior to the experiment, either a wild type (WT) or EPS defective mutant (ΔUDP) of *Pantoea* sp. YR343, a rod-shaped, motile soil bacterium isolated from the rhizosphere of *Populus deltoides*, was grown overnight, diluted, and grown again to an optical density (at 600nm) of 0.1 (OD_600_ = 0.1)^38^. The bacteria was then loaded into the reservoir and seeded into the device for one hour. After one hour, the reservoir was exchanged for pure R2A media, keeping a constant hydrostatic pressure, and the device was imaged hourly for 21 hours. Additional tubing at the outlet carried effluent from the microfluidic platform to dish on a scale where it was weighed every 10 minutes to monitor the flow rate through the platform. Prefilling and sealing the dish minimized evaporation.

Suspending the reservoir 32 cm above the platform resulted in a constant control flow rate of approximately 5 μL/min. After cells were seeded in the platform, this flow rate steadily dropped off over time, approaching 1 μL/min for both strains after 21 hours (Figure 2, a). Despite a reduced capacity to create EPS, a precursor to biofilm development, the ΔUDP strain showed no significant difference in flow rate change compared to the WT strain. However, change in flow rate over time in this heterogeneous porous network was drastically different than the abrupt, catastrophic clogging that other researchers have seen from similar experiments in a single tortuous channel^26^. Presumably, this is because, as bacteria obstruct one preferential flow path, other preferential flow paths open up and divert flow around the clog. This diversion of flow, or remodeling of the porous network, would have environmental implications for chemical transport in natural porous media. Other researchers have found that, as bacteria grow to clog pore spaces, flow redirects to provide nutrients to slower growing colonies^9^. This suggests that, although rapidly dividing bacteria have the advantage in most bulk-media laboratory environments, slow growers may find themselves in advantageous niches within these natural pore networks.

**Figure 2:**
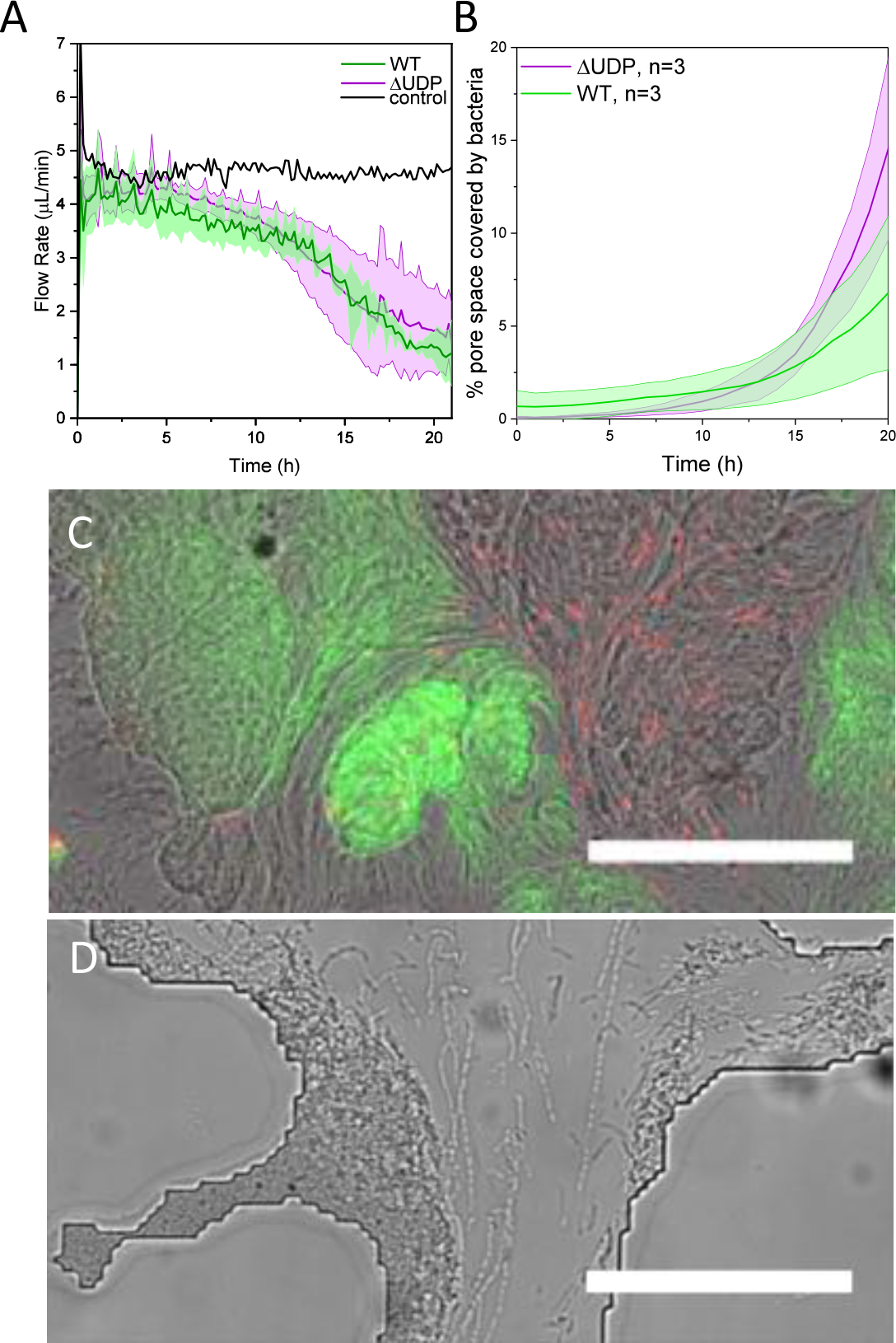
A WT strain and an EPS defective mutant (ΔUDP) grow within the heterogeneous porous media platform. A) Both strains cut off flow at the same rate (n=4, average values shown by dark line while error (standard deviation) is filled with a lighter color) B) The ΔUDP strain grows to cover more area of the device. C) Flow induces a globular phenotype for the WT strain and a D) linked-cell phenotype

Despite both bacterial strains having a similar impact on the overall flow rate through the porous media platform, ΔUDP mutant cells surpassed the WT strain in pore space coverage by the end of the experiment with 10.7% of the pore space covered in live cells versus 5.7% for the WT strain (Figure 2, b). Given that the EPS defective strain cuts off flow at the same rate as the WT, but grows to cover more area, it would seem advantageous for bacteria to produce less EPS under similar flow conditions in the soil. However, this system did not test bacterial competition, antibiotic susceptibility, or challenges to the community that would favor a robust biofilm producing strain^39^.

Hydrodynamic conditions in the pore spaces did alter the phenotype of each strain compared to static laboratory conditions (i.e. agar plate assays). In the pore network, the WT strain formed compact, globular patches of living cells with a diameter approximating the size of the pore in which the bacteria were confined (Figure 2, c). The ΔUDP mutant exhibited phenotypes that depended upon the velocity within the pore space. In pores with flow the ΔUDP cells formed long chains of linked cells, while cells in static pores retained a single cell, rod-shaped structure (Figure 2, d). This ΔUDP phenotype was an important precursor for pore space clogging *in lieu* of EPS formation; strands of living and dead cells spanned multiple particles, forming a foundation on which more cells could attach and grow. The linked-cell phenotype manifested itself across the pore network in a streamer-like distribution of ΔUDP cells that follows the general direction of flow (Figure 3, b). The WT strain, however, exhibited a much patchier distribution of growing microcolonies across the entire pore space (Figure 3, a). A patchy distribution of soil organisms is traditionally thought to be a direct result of heterogeneous resource allocation^40,3^. In this system, nutrients are universally dispersed in the media and continually replenished. However, at later time points, many pores were completely cut-off from flow, and local chemical gradients in nutrients, secondary metabolites, or signaling molecules may be influencing the patchy distribution of WT cells.

**Figure 3:**
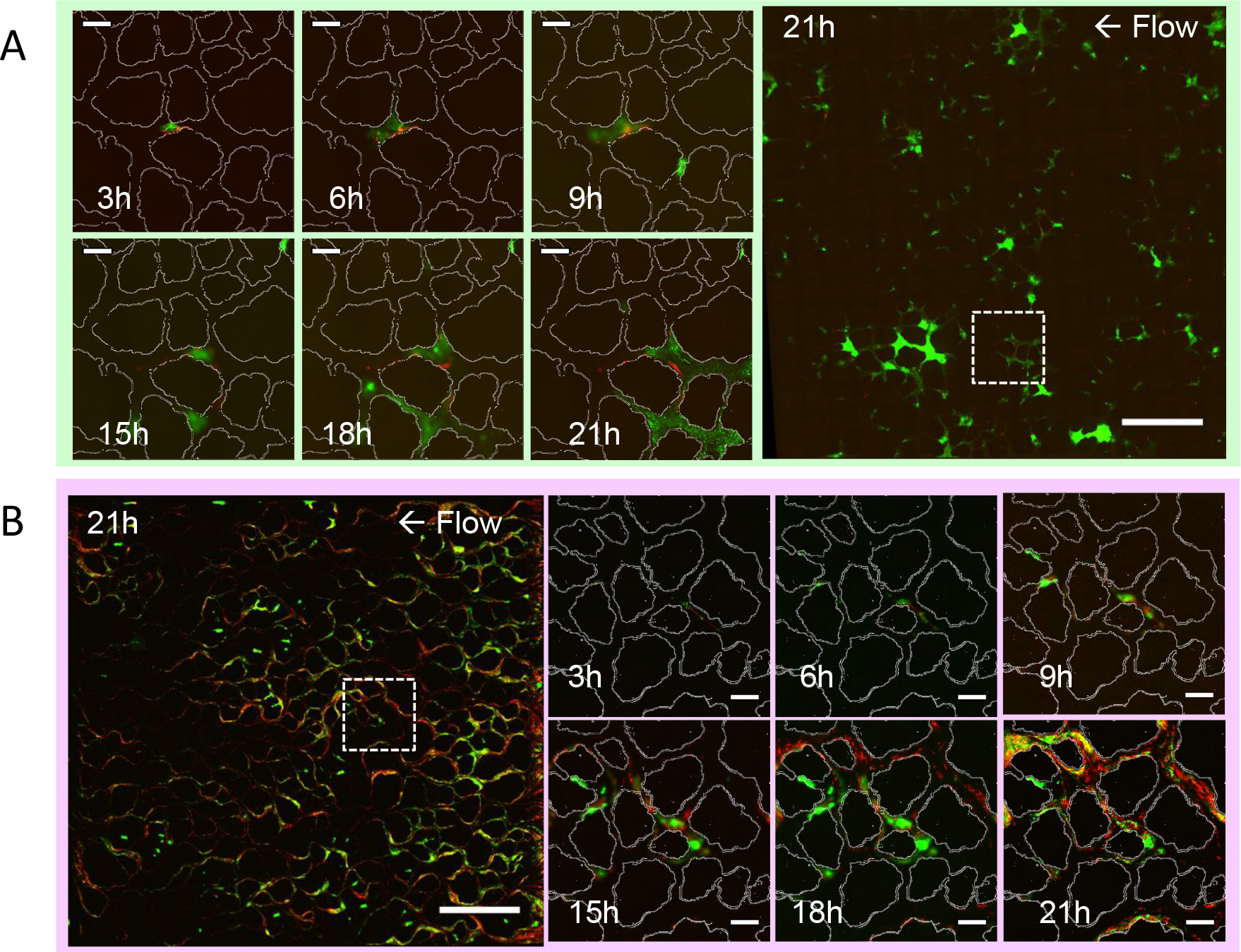
Representative fluorescent images of A) WT and B) ΔUDP cells in the platform at 21 hours (scale bars = 1mm) and in a subset (indicated by dashed box) of the device over time with the particle design overlaid (white) (scale bars = 100 μm)

The spatial distribution of both bacteria strains across the porous network after 21 hours appeared to evolve from the location of the initially attached bacteria. Therefore, we next examined the simulated hydrodynamics within the pores where we saw the highest incidence of initial attachment, or seeding.

### Correlating initial bacteria attachment with flow hydrodynamics

Given the reproducibility of the microfluidic particle layout, one might expect deterministic and predictableseeding within the network and clogging of the same pores through multiple replicate experiments. However, after one hour of seeding, bacterial fluorescence was not always located in the same pores, neither between bacterial treatments (WT or ΔUDP) nor between replicates (n=3). This suggests that, despite the repeatable hydrodynamic conditions and particle layout, bacteria exhibit stochastic, rather than deterministic, seeding in a heterogeneous pore space. Both stochastic and deterministic processes are generally accepted as occurring simultaneously in the formation of local microbial communities^41,42^. While stochastic processes are usually associated with ecological drift, or the relative changes in species abundance due to reproduction and decay, stochasticity is less frequently associated with ecological dispersal or the transport of cells across distances^43^. Dispersal of bacterial species is credited with increasing local diversity and can have dramatic implications for plant-microbiome formation^44^. Our finding that dispersal has a stochastic relationship to pore space hydrodynamics may help inform ecological theories of microbial community formation^45^.

While bacteria did not always initially clog the same pores between replicate experiments, there was a trend in the hydrodynamics of pores where cells were initially captured. In comparing the simulated hydrodynamic conditions of the pores where cells were ‘trapped’, the probability of clog initiation was high for pores with a zero shear rate and increased with the pore space shear rate from 75 s^−1^ up to the maximum shear rate in the device (1000 s^−1^)(Figure 4). In the case of low shear rates (< 75 s^−1^), pore space velocities ranged widely from 0-10,000 μm/s, and for velocities less than bacterial swimming speed (30-75 μm/s for other motile soil bacteria) cellular motility can overcome pore space hydrodynamic conditions so that cells can overcome fluid flow and respond to local chemical signals and nutrient gradients.

**Figure 4:**
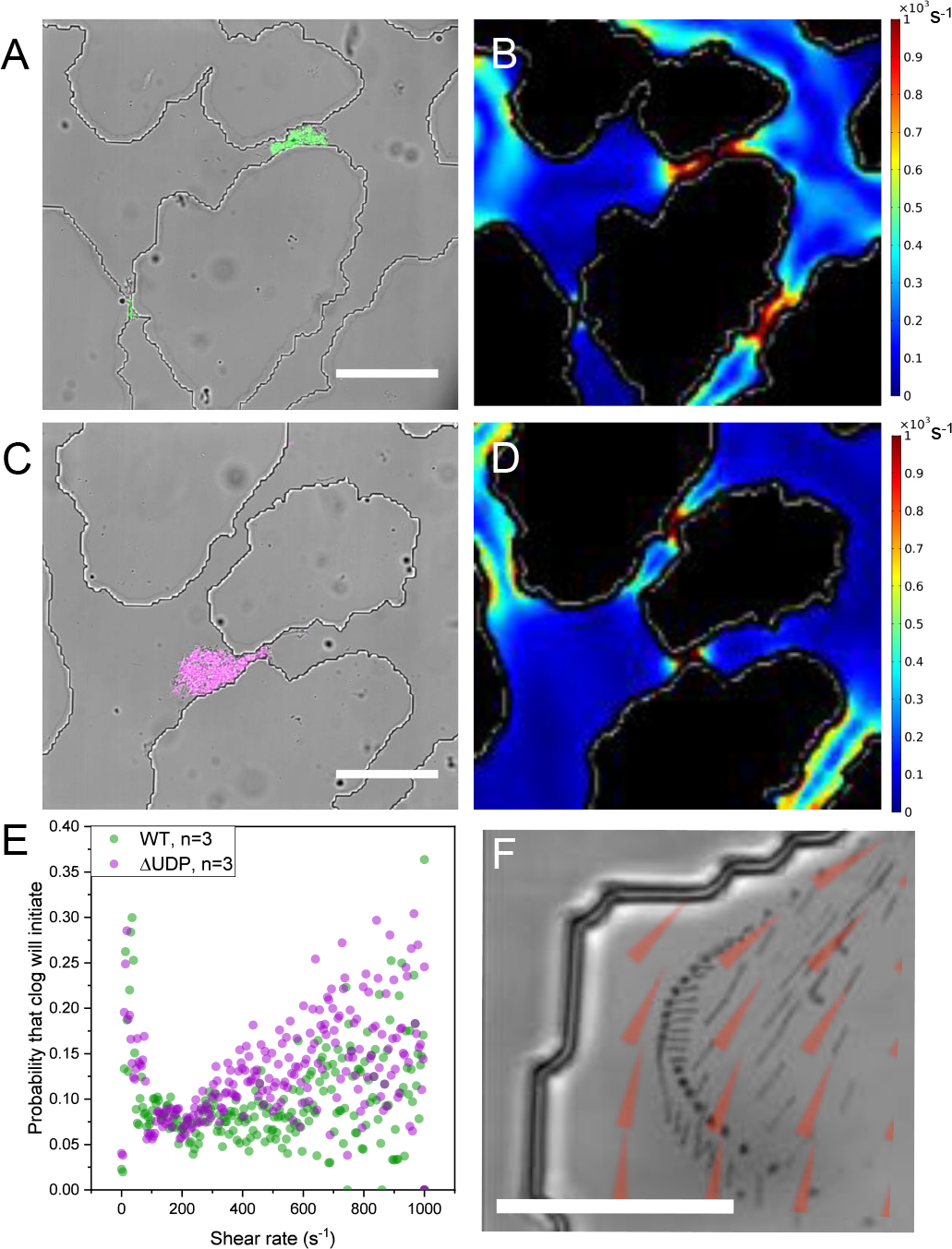
Differential interference contrast images from representative experiments overlaid with fluorescent images of A) WT and C) ΔUDP cells show that the location of initial bacteria seeding correlate to (B,C) areas of high shear from the COMSOL simulations.(scale bars = 100 μm) E) The probability of a clog occurring in a pore is high at zero shear and increases again with shear rate. F) A minimum intensity projection over time of WT bacteria cells (black) moving across the velocity gradient (overlaid red arrows from COMSOL simulation) confirming rheotaxis in the pore spaces. (scale bar = 25 μm)

At high shear rates cells became trapped against particle surfaces by a transport phenomenon known as rheotaxis^46^. In bacteria, rheotaxis is passive directed movement across velocity gradients due to lift forces on the bacteria flagella and drag forces on the bacteria cell body^47^. Because velocity at the surface of a particle is zero, following the no-slip boundary condition, high shear rates tend to occur close to particle surfaces thus encouraging cell-particle interactions and, subsequently, biofilm initiation. Videography of cell trajectories confirmed that rheotaxis was directing cells across velocity gradients toward particle surfaces (Figure 4, f).

An algorithm was created (see Methods section) to quantify the average width of each pore, and this parameter was compared to average shear rate within the same pore (Figure S1). While pore width and shear rate were not directly correlated, the highest shear rates occurred in pores on the smaller end of the size spectrum. This result suggests that soils with small pores (e.g. silt) will capture more bacteria via rheotaxis during initial inoculation than sandy soils. Plants may already be taking advantage of this soil-bacteria interaction. Through growth and mucilage production, roots compact and aggregate soil particles into a “rhizosheath”, thereby improving water retention and effectively reducing pore size, which may increase microbial rheotaxis around the roots and aid in plant microbiome formation^48^.

In addition to being an important factor in trapping bacteria during the initial stages of biofilm seeding within the pore space, we next tried to elucidate the role of rheotaxis in shaping growing biofilms.

### Predicting the spatial evolution of bacterial biofilms

Once bacteria have attached to the surface of a particle, their distribution is affected not only by flow, but also by growth^49,50^. Daughter cells may detach and be carried downstream to attach in another location. They can also reattach upstream when cells have specialized characteristics like curved structure or twitching motility^51,52^. Under these growth conditions, the distribution of live cells and pore scale hydrodynamics are coupled. As resistance in the system builds and overcomes the pressure at the device inlet, pore flow decreases locally and bacteria can use swimming motility as a dominant transport mechanism.

To determine if shear trapping via rheotaxis was dictating the spatial distribution of cells during the growth phase, we converted images of living cells in the device at an initial time (t_i_) to CAD files and incorporated them into the geometry of the COMSOL simulation (Figure S2). The shear profile was then recalculated and compared to the distribution of living cells one hour later (t_i+1_) using object overlay analysis. The probability of co-localization between cells at t_i+1_ and shear simulated from t_i_ was relatively constant for i=3, 9, and 15 hours indicating that shear trapping is no longer a dominate mechanism determining the spatial distribution of a growing biofilm.

Next we conducted a nearest neighbor interaction analysis for bacterial fluorescence in pairs of images (t_i_ and t_(i+1)−i_) for i = 3, 9, and 15 hours^53,54^. The probability of a cell cluster encountering a neighboring cluster peaked at the relatively close distance of 6, 15, and 20 μm for growing cells at i= 3, 9, 15 hours, respectively (Figure 5). This result indicates that the distribution of the growing biofilm over time depends largely on the historical location of the mother cells. By looking at the change in bacterial fluorescence over time, it is apparent that the biofilms grow through pore spaces in a downstream direction.

**Figure 5:**
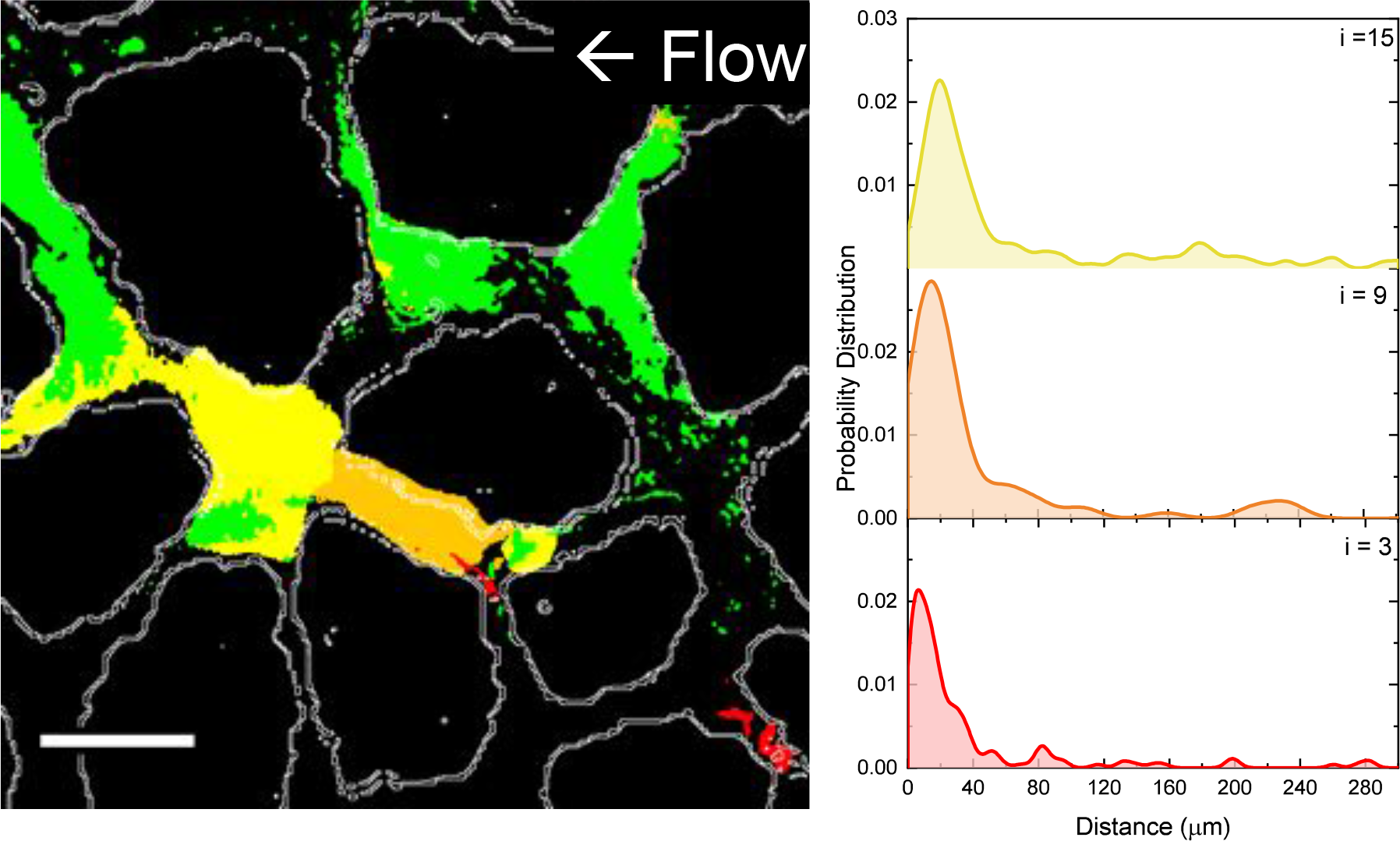
(left) The change in WT bacterial fluorescence over time from 0-3 hours (red), 3-9 hours (orange), 9-15 hours (yellow), and 15-21 hours (green) show that bacteria biofilms are growing steadily in a downstream direction. (right) The nearest neighbour probability for i=3, 9, and 15 hours shows that nearest neighbors are within a 20 μm radius (not considering pore space tortuosity).

Other authors have also determined through nearest neighbor analysis that most soil bacteria interactions occur within a 20μm radius^10^. If the fluid dynamics of this system were static, this result would suggest that bacteria interactions are largely clonal, occuring mostly between descendents of the same ancestor cell. However, flow acts to disperse cells, metabolites, and signalling molecules to downstream microcolonies, increasing the extent of bacterial interactions, albeit in one direction. Therefore, biofilm growth and EPS production, although jeopardizing the incoming flow of nutrients by choking off flow, may provide a mechanism for increasing local communication and bi-directional exchange (e.g. horizontal gene transfer) between bacterial microcolonies in the soil. Results from this work can be incorporated into individual based models of biofilms and used to predict bacterial processes underground such as bioremediation, nutrient cycling, and rhizosphere interactions^55,56^.

The results from this work testify to the ability of heterogeneous matrices to influence emergent microbial properties (i.e. cell phenotype, spatial distribution, and surface attachment) compared to bulk laboratory methods. The use of a faithfully replicated physical environment reveals that microbial dispersal is a stochastic process and particle attachment depends on shear rate within particle spaces. It is imperative that complex systems like these be used to accurately study natural microbial characteristics and community interactions. While this platform only replicates the 2D physical structure of natural sands, increasing complexity can be incorporated into the design, including chemical treatments and nutrient gradients, to recreate natural microbial habitats and emergent bacterial phenomena in a fully defined, parameterized approach.

## Methods

### Device design and fabrication

Using a published granular media algorithm with defined input parameters (Supplementary Table 1), the porous media design was created to replicate the shape distribution of sand particles from Tecate, Mexico^36,37^. The algorithm generated a .bmp file of the particle layout that was vectorized in LayoutEditor to create the CAD design. The particle size of the CAD file, defined by the D_50_ gradation, was scaled to 0.13 mm to reflect smaller sand particles. The design’s porosity (0.38), defined as the ratio of open space to solid space, approximated the porosity of sandy loam (0.25-0.35) ^35^. A bifurcating inlet and outlet were added to the CAD to uniformly distribute bacteria across the design.

The design was transferred onto a chrome mask using a Heidelberg DWL 66 mask writer and replicated onto a silicon wafer using standard photolithography techniques with an NFR photoresist and Bosch etch (Oxford RIE) for to give a final depth of 10μm. This depth was chosen to confine bacteria to a single focal plane during the course of the experiment. Trichloro(1H,1H,2H,2H-perfluoro-n-octyl)silane), 85°C, 60 min) was evaporated onto the surface of the silicon wafer in order to prevent adhesion during the polymer molding process. The microfluidic devices were replicated from the silicon master using poly-dimethylsiloxane (PDMS; 5:1 PDMS base to curing agent, wt/wt; Sylgard 184, Dow Corning) and cured overnight at 70°C ^57^. Inlet and outlet holes were created using a 1.5 mm biopsy punch and the PDMS was bonded to a glass coverslip using air plasma treatment.

### Bacteria Growth

*Pantoea sp*. YR343, a plant growth promoting rhizosphere isolate from the *Populus deltoides* microbiome, was chosen for its relevance to soil ecology and for its rod-shape and flagellar motility^38,58,59^. Fluorescent strains were constructed by integrating GFP into the chromosome of Pantoea sp. YR343 using the pBT270 and pBT277 plasmids as previously described^38^. The EPS mutant was isolated by screening a Pantoea sp. YR343 transposon library on Congo Red plates. The transposon insertion site for the EPS-defective mutant was confirmed to disrupt gene PMI39_01848, which is located at the beginning of an operon with homology to EPS biosynthesis gene clusters found in Pantoea stewartii and Erwinia amylovora^60^.

Fluorescence varieties of the WT and ΔUDP strains were grown overnight prior to the experiment on liquid R2A media (TEKnova, Inc.) at 37°C. Bacteria cultures were diluted 100x in fresh R2A the next morning and allowed to grow again to mid exponential phase at an optical density (600nm) of 0.1, as measured by UV spectroscopy.

### Experimental Setup

Large (for media) and small (for seeding bacteria) reservoirs were created by piercing a hole at the bottom of 50mL and 15mL centrifuge tubes and securing a 23 gauge blunt-tip needle to the tube with polyurethane glue (DUCO cement). Inlet and outlet Tygon tubing was measured to 25cm and 44cm, respectively. The microfluidic device was placed under vacuum for one minute to facilitate pre-filling with R2A media. Once filled, the device, inlet and outlet tubing, and the reservoirs were UV sterilized for 15 minutes in a UV Stratolinker oven. The materials were assembled on the microscope stage before the start of the experiment as shown in Supplementary Figure 3.

A bacteria culture (either WT or ΔUDP strain) was loaded into the smaller reservoir so that the top of the liquid was 32cm above the microfluidic platform inlet. This ensured a consistent pressure-driven flow at 3136 Pa following the hydrostatic pressure head equation (P=ρgh) assuming that the cultures have the same density (ρ) as water. The tubing was attached to the microfluidic inlet and the cells were seeded into the device for 1 hour, after which the reservoir was exchanged for the larger R2A media reservoir taking care to keep the same hydrostatic pressure. Propidium iodide (PI) (0.025μΜ) was added to the R2A reservoir to stain dead cells over time.

Effluent from the microfluidic platform was carried to a Petri dish on an analytical scale. The dish was sealed with Parafilm to limit evaporation and pre-filled with water to quickly establish a steady-state vapor pressure in the dish. The scale was imaged every 10 minutes and the flow rate was calculated with the assumption that the liquid had the same density of water.

After seeding, bacteria within the microfluidic platform were imaged every hour using an inverted microscope (Nikon Ti-Eclipse) with FITC and TRITC epifluorescence to capture both living and dead cells. Image acquisition was carried out with a 20x objective using Nikon Elements software and a back-illuminated iXonEM+ 897 camera (Andor).

### COMSOL Simulation

The porous media CAD was incorporated as the pore space geometry in a 2D laminar flow (spf) COMSOL Multiphysics (version 5.2) simulation. The fluid space was assigned the material properties of water while the particle space was subtracted from the geometry. One side of the design was assigned as an inlet boundary with an entrance pressure of 313.6 Pa (3,136 Pa calculated from the hydrostatic pressure divided by 10 μm for the device height) and the other side of the design was assigned an outlet pressure condition of zero Pa. Due to anisotrophy in particle layout, care was taken during experimental setup to match the experimental inlet with the simulated inlet. An extra-course free triangular mesh was used to partition the geometry into finite elements. The laminar flow study used a stationary GMRES solver (50 iterations) with a relative tolerance of 0.001.

An outlet flow rate from the simulation was calculated by multiplying the line integrated velocity magnitude at the outlet by the device depth (10 μm). This value (4.706μL/min) closely matched the experimental control flow rate of only R2A media of 4.48μL/min (n=1). Experimental particle image velocimetry using fluorescent 2μm carboxylated polystyrene beads confirmed that the simulated velocity matched experimental average speeds in the pore spaces.

For flow simulations with the biofilms, a fluorescent image of both live and dead cells at a given time (t_i_) was converted to a CAD file (.dxf) and subtracted from the COMSOL geometry. This method worked reasonable well at earlier time points (i=3, simulated flow rate= 4.56μL/min and experimental flow rate = 4.64μL/min) but the experimental and simulated flow rates diverged at i=9 (simulated =4.57μL/min, experimental=3.91 μL/min). This method of modeling biofilms, is therefore limited, and might be improved by adding polymeric material properties to the biofilm geometry and incorporating a solid-liquid interaction into the simulation. However, the rheological properties of this particular bacterial biofilm remain uncharacterized.

### Image Analysis

To remove out-of-focus autofluorescence from debris and background noise, images were processed using Fiji^61^. For each image, the background fluorescence was removed (sliding parabloid, 700μm) and then images were manually thresholded to further reduce noise. Total fluorescence area was analyzed using the particle analyzer feature.

#### Object Overlay Analyses

For object overlay analyses with the COMSOL simulations, experimental images were rotated, cropped, and downscaled to align with the same region of interest in the COMSOL shear rate profiles. A composite image was created from the experimental and COMSOL images using the AND Boolean operation. The histogram of greyscale pixel values from this composite image provided information on the shear rate values that were co-localized with each bacterial fluorescence pixel. To obtain the “probability that a clog will initiate” metric for each shear rate value, the shear rate associated with each bacterial fluorescence pixel at i=1 was normalized by the total number of pixels from the COMSOL simulation with the same shear rate value.

#### Nearest Neighbour Interaction Analyses

Change in bacterial fluorescence was calculated by subtracting an image at an initial time (t_i_) from an image one hour later (t_i+1_). The resulting image (t_(i+1)−i_) was compared to t_i_ using the Mosiaic Interaction Analysis Plugin to determine nearest neighbours for i=3,9, and 15^54^.

### Pore Space Characterization

A pore characterization program was created as an addendum to the 2D granular space simulation (GSS) reported by Mollon and Zhao^37^. The purpose of the complementary pore characterization program was to quantify the average width of each pore generated using the GSS. Pores are represented as linear segments in the GSS and the length of the linear segment is defined here as (L). The blue linear network shown in Figure S1 (top panel) represents the Voronoi tessellation. An iterative algorithm is required to converge to an approximation of the real tortuous pore pathlength. The pore width is ultimately derived from this pore pathway reconstruction. Figure S1 shows in the initial distribution of transverse pore lines in red 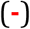 while the final transverse profile representing an approximation of the tortuous pore pathlength is shown by the population of black lines (**-**). Details of the pore characterization program can be found in the Supplementary Information.

## Acknowledgements

This work was supported in part by the Genomic Science Program, U.S. Department of Energy, Office of Science, Biological and Environmental Research, as part of the Plant Microbe Interfaces Scientific Focus Area (http://pmi.ornl.gov). The fabrication of the microfluidic platforms was carried out in the Nanofabrication Research Laboratory at the Center for Nanophase Materials Sciences, which is a DOE Office of Science User Facility. JAA is supported by an NSF graduate research fellowship DGE-1452154

## Supplementary Figures and Information

**Figure S1:**
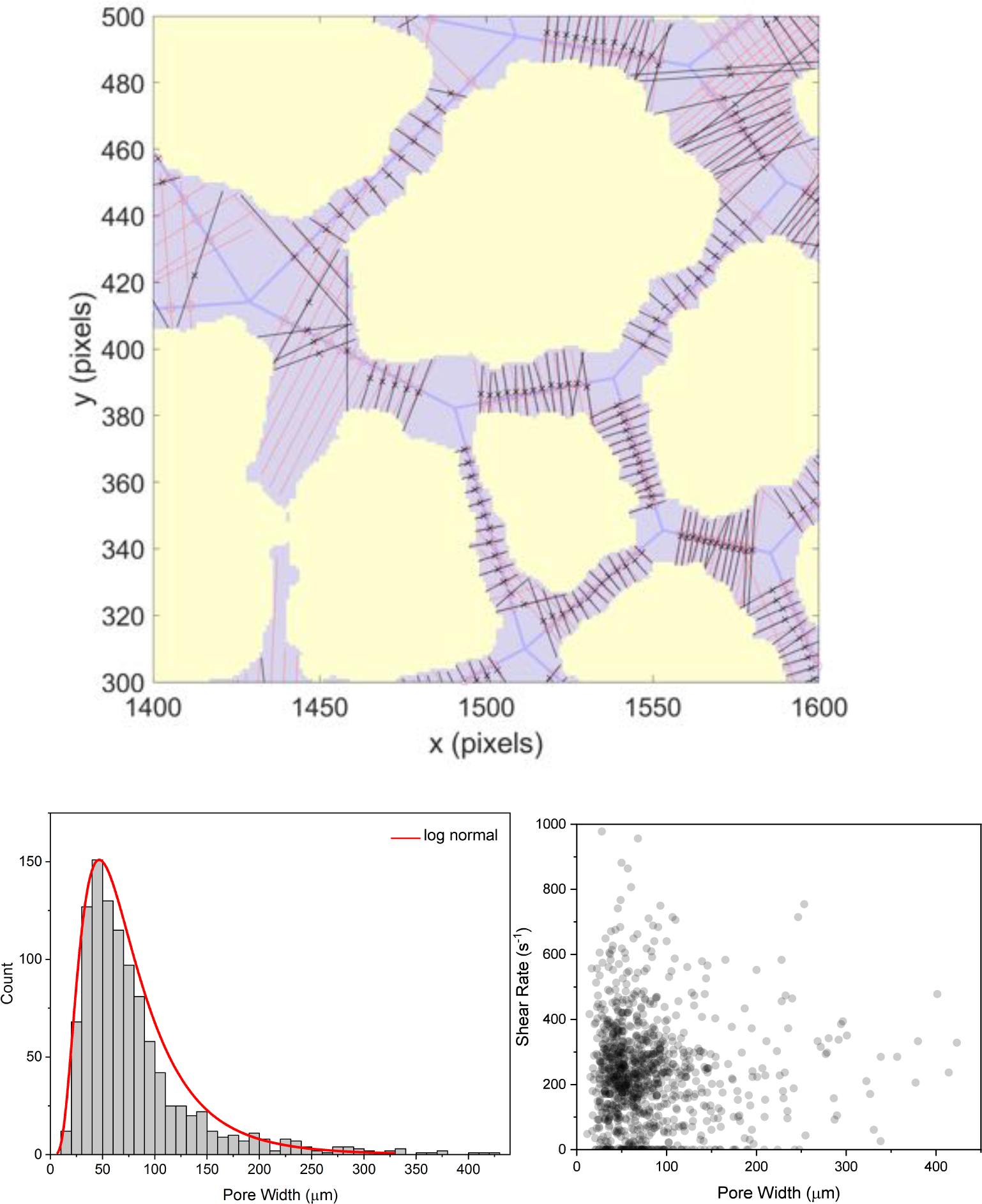
(top) Pore characterization program results shown in a 200 pixel × 200 pixel region–of–interest sampled from a larger 2D granular space of roughly 2000 pixels × 2000 pixels. Pores in the tessellation pattern with total path length less than 10 pixels were not included in the pore characterization statistics. (bottom left) A histogram of the pore widths shows a close resemblance to a lognormal function. (bottom right) Every pore is plotted by its average shear rate and width.

**Figure S2:**
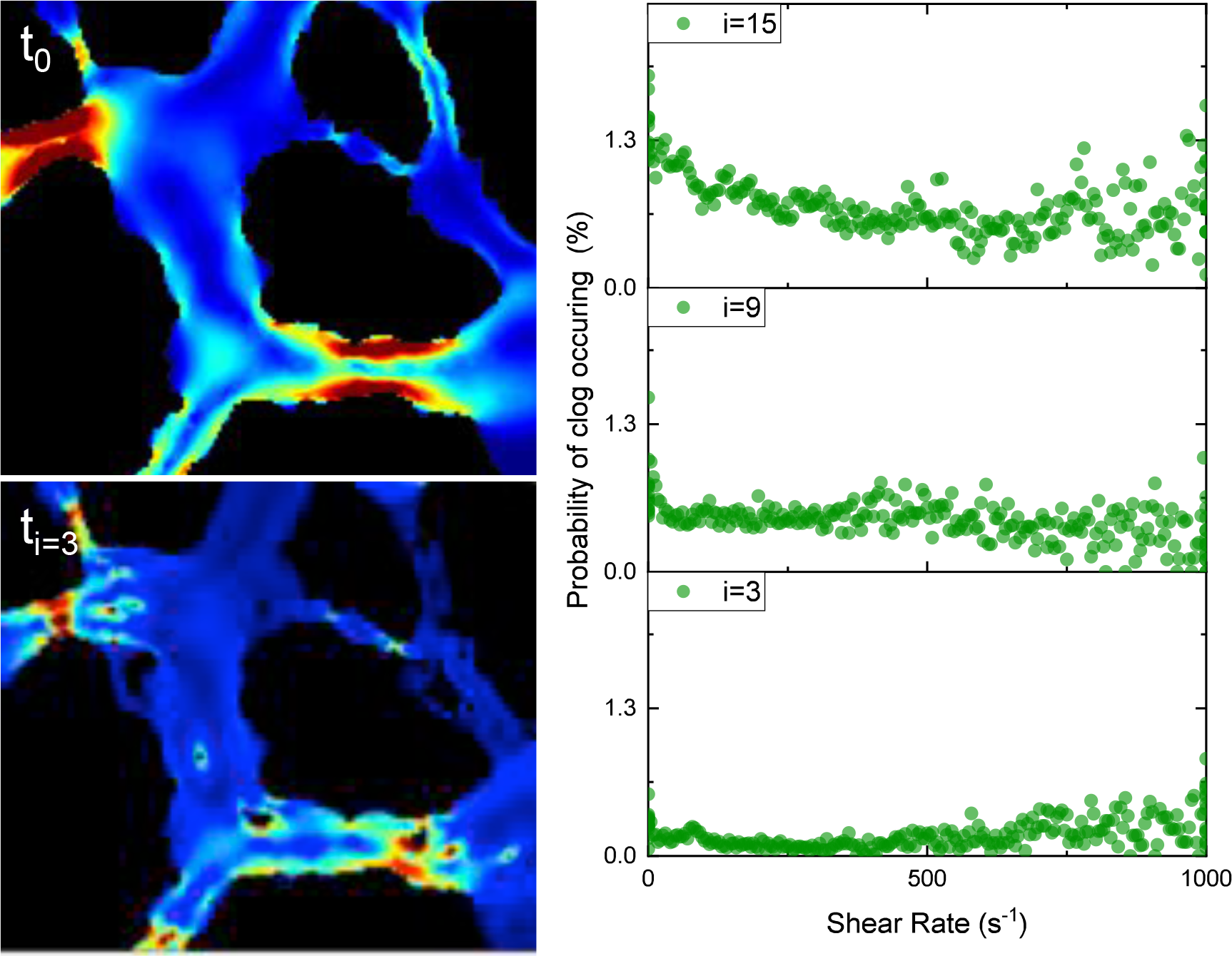
(left) An example of the simulated shear rate before (t_0_) and after subtracting the biofilm from the geometry (t_i=3_) shows that biofilm influences pore space shear rates. (right) The relationship between shear rate and the probability of a clog occurring is relatively constant over time.

**Table S1:** Input Parameters for the particle generating algorithm^37^.

|  |  |
| --- | --- |
| **Number of Particles** | **500** |
| **SurfaceCOV** | 0.8 |
| **Orientation** | 0 |
| **Anisotropy** | 1 |
| **itermax** | 1.00E+06 |
| **error** | 0.002 |
| **solid fraction** | 0.65 |
| **nvar0optim** | 6 |
| **covspectrum** | 0 |
| **typespectrum** | 1 |
| **file spectrum** | tecate |
| Dmax | 0.02 |
| Rmin | 0.02 |
| pmax | 0.05 |

**Figure S3:**
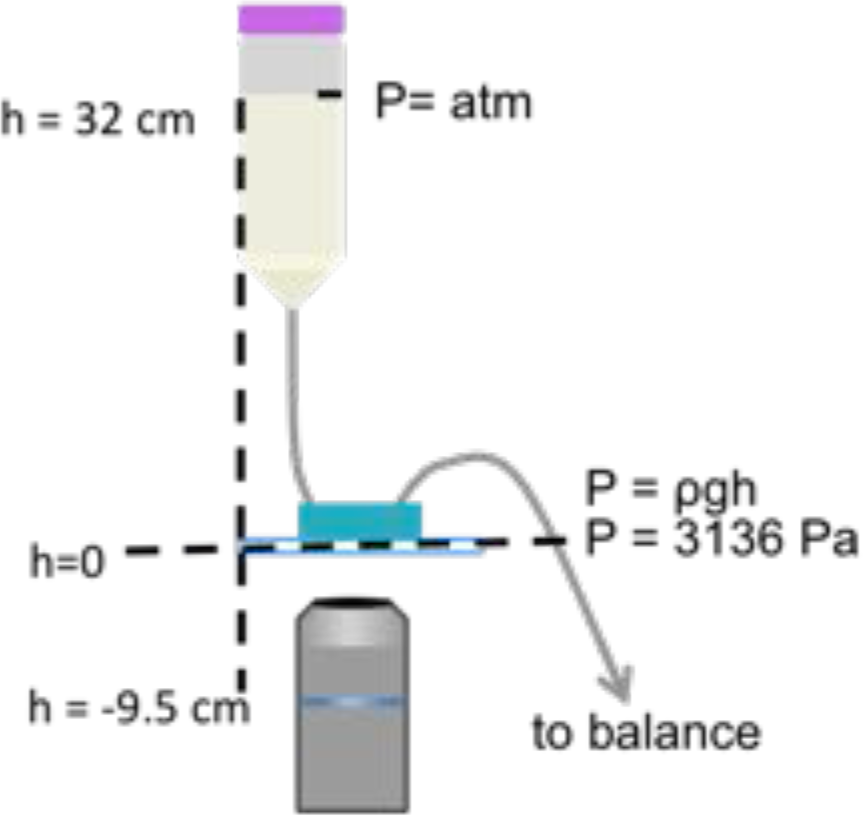
A schematic illustrating the experimental setup and the calculation of hydrostatic pressure. The lid on the reservoir tube was loosely fitted to assure atmospheric pressure above the liquid.

## Supplementary Information

### Pore characterization program

#### Purpose

The pore characterization program was created as an addendum to the 2D granular space simulation (GSS) reported by Mollon and Zhao^37^. The purpose of the complementary pore characterization program is to quantify the statistical attributes of each pore generated using the GSS. Pores are represented as linear segments in the GSS and result from a Voronoi tessellation^37^. The length of the linear segment is defined here as (L). The blue and linear network shown in figure S1a represents the Voronoi tessellation. Additional pore attributes of interest that are not provided by the GSS include the average pore width, the variation of the pore width and an approximation of the actual tortuous pathlength between neighboring grains. The pore characterization program is now summarized.

#### Pore Definition

The pore characterization program is described from the point–of–view of a single pore for clarity. The linear segment defining the pore, derived in the GSS, is sub–divided into multiple, equally spaced nodes (M). The total number of sub–division nodes (M) is an input parameter and ultimately controls the spatial sampling resolution of the pore. Each individual sub–division node is referenced using (m). The user also provides a parameter, range 0–1, that represents the fraction of segment ignored during segment sub–division, defined as (f). The purpose of this parameter is to avoid sampling the pore width at the ends of the segment where the pore is poorly defined in space, as described next.

Multiple pores converge at a common node due to the Voronoi tessellation used to create the granular space using GSS. In combination with the GSS cell filling procedure, these nodal areas tend to produce relatively large open spaces at pore entry and exit surfaces. During the sub–division procedure in the pore characterization program, nodes introduced near the segment ends sample these open spaces, instead of the pore space, which leads to an overestimate of both the mean pore width and mean pore width variation. To avoid this effect, the parameter (f) was introduced to prevent nodal points near the segment ends. This is accomplished by leaving a linear space equal to (f*L/2) free of sub–division nodes at the pore entry and exit. Quantitatively, this results in the following nodal range of points (**P**);

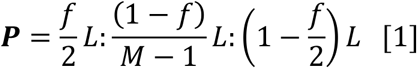

where **P** is defined along the linear segment. The origin is assigned arbitrarily and can be either segment node. In this work, the linear segment node with the smallest (x) value is taken as the origin. This arbitrary designation was made simply because the general fluid flow direction during experiments is oriented along (+x). The pore is therefore initially defined by the vector;

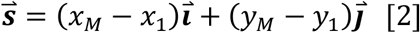

The distribution of initial nodes **P** are indicated with open red circle data points 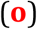 in figure S1a.

#### Pore Shape Extraction

An iterative algorithm is required to converge to an approximation of the real tortuous pore pathlength. The pore width is ultimately derived from this pore pathway reconstruction. *Convergence is obtained when the following two conditions are met for each sub–division node along the pore; (1) the center position (x,y) of the pore node (m) has been found and (2) this center position (x,y) also intersects the line spanning the pore which best aligns with the surface normal on each side of the pore*. The user defines the number of iterations to convergence. Figure S1a shows the outcome of an acceptable level of convergence for pore definition using the pore characterization program. Specifically, the black data points (**x**) show the final distribution of pore sub–division nodes following convergence. A summary of the iterative algorithm is now provided.

### The Iterative Solver

The initial condition is a series of equally spaced nodes lying on the linear segment defining the pore (*see previous section*). The normal vectors, oriented orthogonal to the initial pore vector 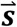, are calculated for each node defined in the range **P**. This calculation requires that the binary, digital image of the granular space map be superimposed with the pore network defined by the Voronoi tessellation. Ultimately, this overlay is necessary for determining the pore width at each sub– division node. In the binary digital image, a pixel value of 0 represents occupied granular space while a pixel value of 1 signifies open ‘pore’ space. The pore width per node (m) is calculated in the following way.

First, the normal vector per node is derived from the local tangent vector to the current pore path estimate. *Importantly, this tangent vector is calculated using the two nearest neighbor nodes as*;

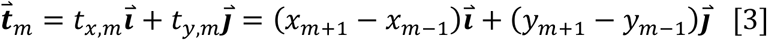

*where the index (m) represents the current node of interest*. By using the 1^st^ nearest neighbor nodes, the solution propagates in an interconnected way through the pore giving a smooth estimate of the tortuous pore pathlength at convergence. The normal vector 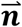 is derived using;

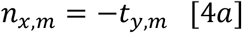

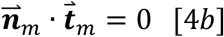

yielding the components n_x,m_ and n_y,m_.

Next, a line is cast through the node (m) aligned along 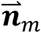. An estimate of the pore cross–section at (m) is measured as the displacement (Dx, Dy) between the two points of intersection between the extended line and the neighboring grains forming the pore. Please note, the exact path of the line through each pixel is measured. This minimizes the error incurred when deriving a linear path measurement from a pixelized image to a value of roughly twice the pixel edge length. The current node (m) position (x_m_,y_m_) is then updated as the average value of the two intersection points. This summary represents the complete sequence of events during a single iteration. The process repeats until convergence is reached. Figure S1a shows in the initial distribution of transverse pore lines in red 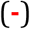 while the final transverse profile representing an approximation of the tortuous pore pathlength is shown by the population of black lines (**-**). For Figure S1a, the number of convergence iterations was set to 40, the number of sub–division nodes per pore was M = 13, and the fraction of total pore length ignored during sampling was f = 0.34. Pores in the tessellation pattern with a total pathlength less than 10 pixels were not included in the pore characterization statistics.

#### Generated Data

Following successful program execution, a text file is saved that contains the following information for each sub–division per pore (m); the pore width, the pore node position and the normal vector defining the pore cross–section at (m). Moreover, this data is provided for each iteration. This information is also provided for all pores in the 2D granular space.

#### Additional Features

Lastly, it is worth briefly mentioning additional features of the program. A program component was included which prevents pores on the simulation boundary from contributing to the final pore characterization statistics. Also, an additional tool is included making it possible to ignore ‘pores’ that are too short in pathlength to represent an actual pore but were nonetheless generated as part of the Voronoi tessellation procedure. These ‘imposter’ pores were typically created when a relatively large number of pores converged at a single vertex in the Voronoi tessellation pattern.

